# Radiation necrosis after radiation therapy treatment of brain metastases: A computational approach

**DOI:** 10.1101/2023.08.01.551411

**Authors:** Beatriz Ocaña-Tienda, Odelaisy León-Triana, Julián Pérez-Beteta, Víctor M. Pérez-García

**Author notes:** These authors contributed equally to this work (Beatriz Ocaña-Tienda, Odelaisy León-Triana).

## Abstract

Metastasis is the process through which cancer cells break away from a primary tumor, travel through the blood or lymph system, and form new tumors in distant tissues. One of the preferred sites for metastatic dissemination is the brain, affecting more than 20% of all cancer patients. This figure is increasing steadily due to improvements in treatments of primary tumors. Stereotactic radiosurgery (SRS) is one of the main treatment options for patients with a small or moderate number of brain metastases (BMs). A frequent adverse event of SRS is radiation necrosis (RN), an inflammatory condition caused by late normal tissue cell death. A major diagnostic problem is that RNs are difficult to distinguish from BM recurrences, due to their similarities on standard magnetic resonance images (MRIs). However, this distinction is key to choosing the best therapeutic approach since RNs resolve often without further interventions, while relapsing BMs may require open brain surgery. Recent research has shown that RNs have a faster growth dynamics than recurrent BMs, providing a way to differentiate the two entities, but no mechanistic explanation has been provided for those observations.

In this study, computational frameworks were developed based on mathematical models of increasing complexity, providing mechanistic explanations for the differential growth dynamics of BMs relapse versus RN events and explaining the observed clinical phenomenology. Simulated tumor relapses were found to have growth exponents substantially smaller than the group in which there was inflammation due to damage induced by SRS to normal brain tissue adjacent to the BMs, thus leading to RN. ROC curves with the synthetic data had an optimal threshold that maximized the sensitivity and specificity values for a growth exponent *β*_∗_ = 1.05, very close to that observed in patient datasets.

**Author summary:** After treatment of brain metastases with radiation therapy, a fraction of patients experience tumor recurrences and others display radiation necrosis (RN). Clinical data shows that the growth of RNs is faster, as measured by the growth exponent, than that of recurrent brain metastases. This reflects the inflammatory nature of the former, and provides a method to distinguish RN in the clinics from relapsing metastatic lesions. A simple mathematical model for the inflammatory response and a more sophisticate discrete stochastic simulator with many biological details were been developed to provide a mechanistic explanation of the differential dynamics of tumor growth versus inflammatory responses after stereotactic radiation surgery of metastatic brain lesions.

## Introduction

Brain metastases (BM) are the most common intracranial malignancies in adults, with around 25% of patients with cancer developing brain metastases during the course of their diseases [1, 2]. Most BMs correspond to primary cancers from lung, breast or from melanoma [2]. Improvements in detection and primary tumor treatment leading to longer survival of cancer patients have resulted in an increase in the incidence of BMs in the last years [3].

Stereotactic radiosurgery (SRS), a form of external beam radiation therapy, is the preferred treatment option for the management of BMs in patients with less than five lesions because of its excellent local control rates [4].

Radiation necrosis (RN) is an inflammatory reaction that appears between 6 and 24 months following SRS in 5% to 25% of treated patients [5]. It appears as contrast-enhancing lesions, resembling the appearance of tumor progressions on routine magnetic resonance images (MRIs) what poses a critical problem for physicians. RN may resolve spontaneously, and does not require further work-up, while progressive BMs require a prompt therapeutic action, typically in the form of open brain surgery. Thus the distinction between the two conditions is a problem with direct clinical implications.

There are two main pathophysiological theories for the development of RN in the brain, 18 and it seems likely the combination of both is the best explanation for the process. One 19 explanation suggests that radiation damages different types of healthy brain cells, 20 resulting in apoptosis [6], while another theory suggests that radiation damages the vasculature [7]. Damage to endothelial cells, which causes cell death in a delayed manner [8] and the release of inflammatory mediators [9], is a critical factor in the emergence of RN. Astrocytes, oligodendrocytes, and oligodendrocyte progenitor cells are also damaged by radiation [10, 11] resulting in the release of hypoxia-inducible factor 1*α* and VEGF [12]. Finally, VEGF induces ICAM-1, increasing the inflammatory response and edema, together with the enhancement seen on CE images.

Although biopsy sampling is considered the gold standard for distinguishing between tumor progression and radiation necrosis [4], there are limitations in its use. Conventional MRI is not appropriate for discriminating between both conditions since they share similar characteristics. Several alternative methods have been employed to differentiate tumor progression and RN, including perfusion MR imaging [13], positron emission tomography [14], the use of artificial intelligence [15] and radiomic methods [16, 17], or changes in anatomical boundaries [18]. However, many of these methods lack independent validation, require complex computational methodologies, or 35 are in need of further study. Consequently, there is a need for diagnostic techniques based on routinely-used imaging methods, such as post-contrast T1-weighted images (T1-WI) [18].

Mechanistic mathematical models have been used to study the growth of untreated metastasis in different scenarios [19–21], the interaction between primary and secondary tumors [22], cancer metastasis networks [23–25], and, and, in recent years, the response to treatment [26–32]. However only a few of them have studied specifically brain metastases [21, 26, 27, 31, 32].

Interestingly, a simplified version of the Von Bertalaffny growth equation fitted using observational longitudinal volumetric patient’s lesion growth data has been recently found [34] to be useful in discriminating both conditions.

The aim of this study was to understand the longitudinal dynamics of RN development and how different are its growth patterns from those of progressive lesions. To do so, two mathematical models were built. The first, a simple compartmental model, allowed the growth of RN events to be understood. The second, based on a discrete stochastic simulator, allowed the response to SRS to be studied, including more details of the biological processes behind post-SRS inflammation.

## Materials and methods

In the planning of radiation therapy and despite improvements in the accuracy of radiation therapy treatments, low doses of radiation are administered to the tissue surrounding the lesion to ensure that not only visible tumor cells but also those that are not visible in medical images are eliminated, to reduce the likelihood of tumor recurrence. However, this approach has the unavoidable consequence of damaging healthy tissues around the metastatic lesion.

While radiation kills tumor cells and reduces tumor volume, it also affects healthy cells, which die by mitotic catastrophe when they are trying to renew. Renewal time for tumor cells is fast, whereas brain cells take months to reach the renewal point. This damage to healthy cells and the subsequent inflammation process governed by the immune system can lead to an apparent increase in tumor volume, which is typically observed several months or even years after SRS [37].

### Compartmental mathematical model

To simulate the development of radiation necrosis over time, a simple mathematical model was first developed, describing the evolution of cells taking into account the impact of SRS on both the tumor cells and the healthy cells in the tissue surrounding the lesion, as illustrated in Fig. 2.

**Fig 1.**
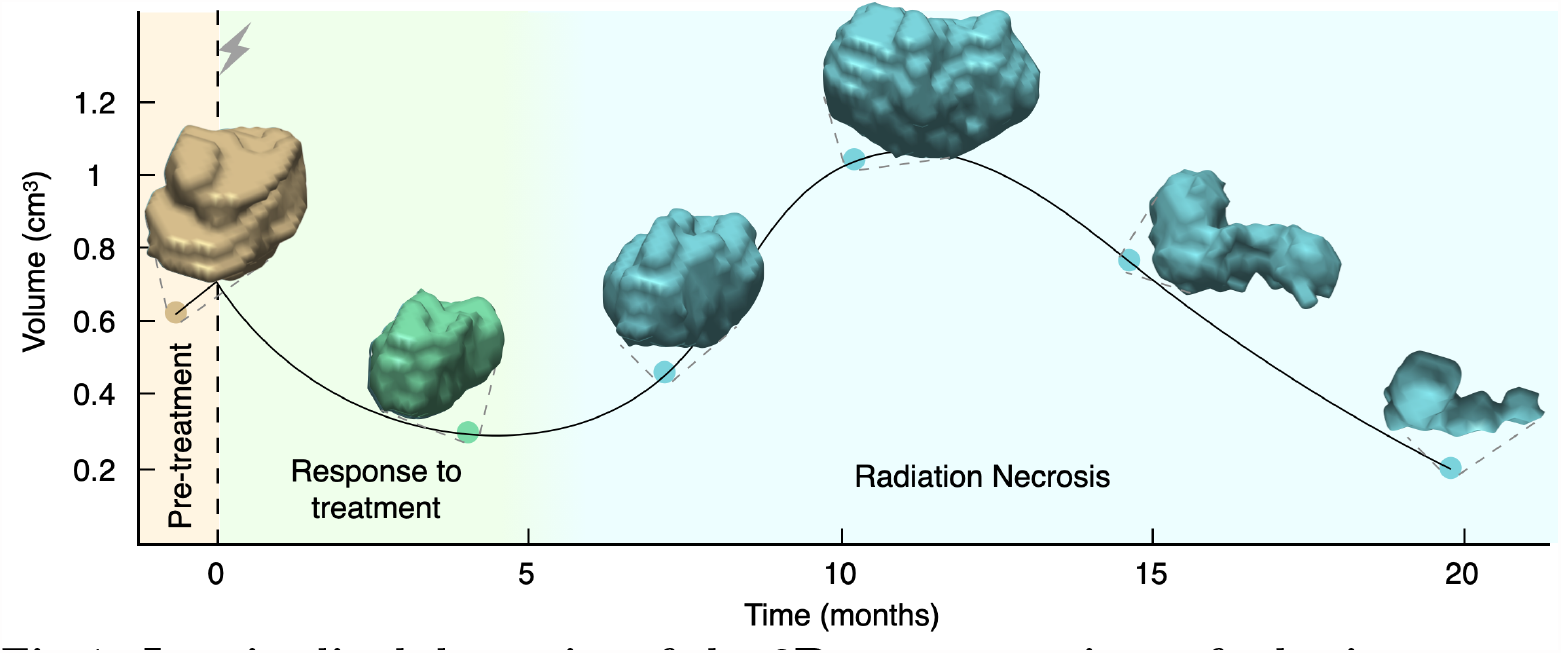
Longitudinal dynamics of the 3D reconstructions of a brain metastasis treated with SRS showing radiation necrosis. Dots correspond to the measured volumes together with their 3D reconstructions and the solid line is the result of interpolating longitudinal volumetric data (shown only to guide the eye). The patient was a 50-year-old male with a non-small-cell lung cancer primary, who underwent a single session SRS with a dose of 20 Gy. In this case, the inflammatory lesion exhibited its peak volumetric expansion around 12 months after treatment. The solid line is a cubic spline interpolation shown to guide the eye.

**Fig 2.**
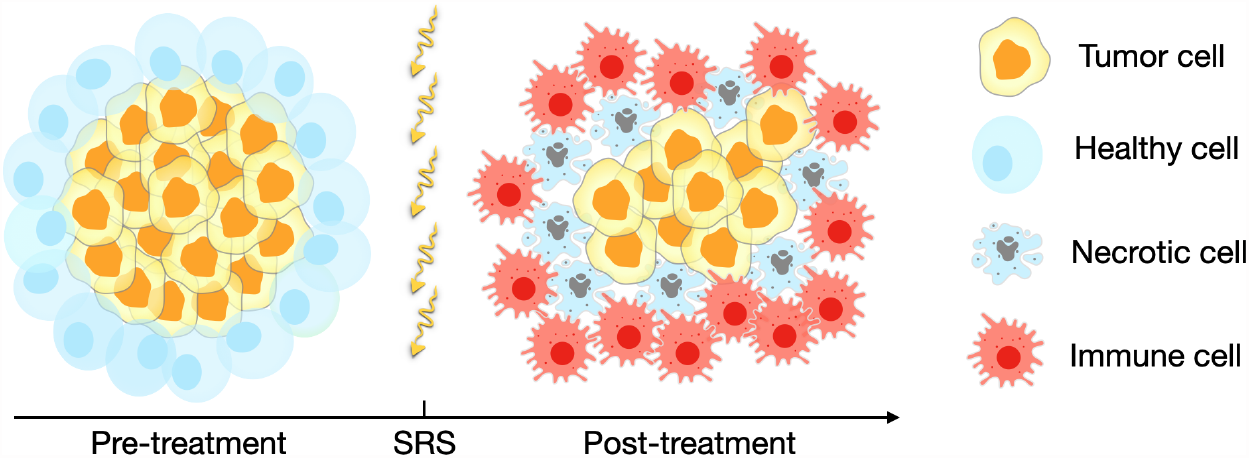
Schematic representation of the populations involved in the response to SRS. The post-treatment scenario represents the growth of radiation necrosis events as defined in Eqs. (1). After SRS healthy cells around the tumor become damaged and die by necrotic catastrophe. The appearance of necrosis stimulates immune cells leading to an apparent growth of the lesion.

It was assumed that the dynamics of the three leading populations involved in radiation necrosis events, tumor cells *T*(*t*), necrotic cells *N*(*t*) and immune cells *I*(*t*) are governed by the following equations:

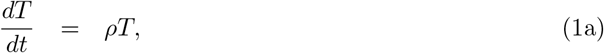

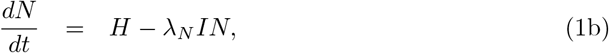

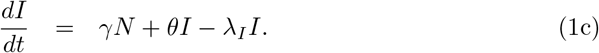

Equation (1a) describes the growth of the tumor cells, which is assumed to be exponential.

The first term in (1b) accounts for the dynamics of healthy cells that are killed by the late effects of radiation when trying to renew as given by

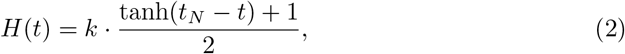

which accounts for the fact that cells have the same probability of reaching the renewal point during a given time period *t*_*N*_. The value of *k* provides information about the number of healthy cells that die at each time. This means that the total number of cells killed by radiation is given by

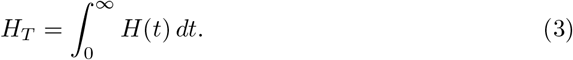

*H* will depend on the initial BM volume, since typically, a region of 1 mm width surrounding the tumor is irradiated to kill potentially infiltrating tumor cells. This is the typical irradiation margin used in BM radiation therapy treatments. The second term in (1b) represents necrotic cells cleared away by immune cells *I*(*t*). Eq. (1c) assumes that the immune response is stimulated by the presence of necrotic cells, while they have a growth rate *θ* and a death rate *λ*_*I*_.

#### Parameters estimation for model Eqs. (1)

The growth rate *ρ* was computed for each lesion from the volumes in the first two points at which growth was observed before any treatment was applied, i.e.

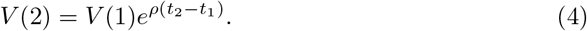

This choice implicitly assumes that the tumor preserves the same growth dynamics after treatment, an assumption taken here for simplicity, although the real dynamics would be more complicated [36].

The death rate *λ*_*I*_ was taken to be 0.07 days^−1^, leading to a half-life of around two weeks, that is the typical lifetime of activated effector immune cells [38, 39].

Parameters *λ*_*N*_, *γ* and *θ* were estimated by fitting the model to available longitudinal volumetric data in [40]. The data consisted of three volumetric time points for each BM, displaying longitudinal volumetric growth. The MATLAB functions `fminsearch` and `ode45` were used to perform the fitting, returning values in the range [1.85 − 2.36] · 10^−11^ days^−1^, [1.84 − 2.06] · 10^−7^ days^−1^ and [0.139 − 0.249] days^−1^, for *λ*_*N*_, *γ* and *θ*, respectively. Thus, the parameter values obtained were very consistent for the different patients. The immune growth rate *θ* was found to be about 5 days, which seems a reasonable value. In contrast, the necrotic cell elimination rate *λ*_*N*_ and the rate of activation of immune cells *γ* were found to be very long. This finding may be influenced by the delayed effect of healthy cell death and the activation of the immune system associated with the appearance of RN.

#### Calculation of the *β* exponent

To compute *β* for the simulations of model (1), the simulated time interval was split into three time slots, and for each slot, a random time value was chosen. This method was designed to mimic the clinical imaging follow-up, where scans are typically performed in three-months time windows but with a substantial variability due to real-world constraints. Those times *t*_0_, *t*_1_, *t*_2_ and the computed volumes *V*_0_, *V*_1_, *V*_2_ were used to compute the growth exponent *β* using the simplified Von Bertalanffy growth equation [33]

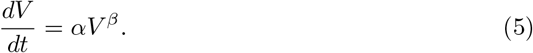

Solving Eq. (5), gives

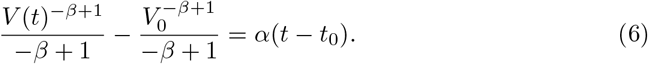

Since there is information on the dynamics at three-time points (*t*_0_, *V*_0_), (*t*_1_, *V*_1_) and (*t*_2_, *V*_2_), the two parameters *α* and *β* can be completely determined by evaluating (6) at the times *t*_0_, *t*_1_, *t*_2_, giving

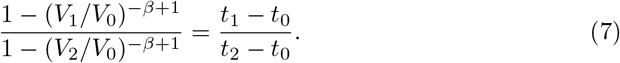

Eq. (7) is an algebraic equation for *β* that was solved by using the MATLAB function fzero (which returns the root of a nonlinear function) for each set of known values *V*_0_, *V*_1_, *V*_2_, *t*_0_, *t*_1_, *t*_2_.

A dataset of 500 virtual BMs was generated for each choice of initial volume, ranging from 0.5 cm^3^ to 3.0 cm^3^. The parameters *λ*_*N*_, *γ*, and *θ* were allowed to take uniformly distributed random values within ranges chosen to include the parameters obtained when fitting volumetric data. Specifically, *λ*_*N*_ was assigned a value within the range [1.8 − 2.4] · 10^−11^ days^−1^, *γ* within [1.8 − 2.1] · 10^−7^ days^−1^, and *θ* within [0.14 − 0.25] days^−1^. By varying the initial volumes and parameter values, it was possible to obtain a wide range of *β* values and to analyze the resulting trends.

### Discrete stochastic brain metastasis simulator (DSBMS)

The mathematical model used for the prior analysis included only a limited number of relevant biological elements. A fuller study was carried out using a discrete stochastic brain metastasis simulator (DSBMS) based on Ref. [41]. The model included six types of cellular population: normal and damaged healthy cells, normal and damaged tumor cells, immune cells and necrotic cells. For these populations, the main biological processes were implemented at the cellular level: mitosis, migration and cell death.

The discrete model focuses on describing cell populations instead of individual cells. The spatial domain was set as a 3D grid discretized in cubic compartments (voxels) of side length Δ*x*, fixed at 1 mm. Each voxel has a specific dynamic that depends on its occupation and surroundings and can contain cells from each subpopulation with an upper limit indicated as local carrying capacity *K*. The different cell populations attempt to perform all basic processes at each time step. These processes can be described by a binomial distribution with a probability associated with the process. That is, the number of cells successfully undergoing division, death, migration or transition to another population within the discrete model are calculated voxel-wise and state-wise at each time step. These numbers are thus calculated by randomly sampling the corresponding binomial distribution, whose *N* will be the number of cells in each population within a voxel, and whose probability will be the rate of the process modulated by the time step length Δ*x*. All processes, their probabilities and associated binomial distributions are described below in detail.

#### Tumor growth in the discrete model

To simulate the growth of the metastatic lesion before SRS, the biological processes of cell division, death and migration were implemented similarly to the model in [41]. Let *n*_*t*_, *n*_*n*_ and *n*_*h*_ be the total numbers of tumor, necrotic and healthy cells inside a given voxel, respectively. The total numbers of proliferating and dying tumor cells are drawn from the binomial distributions 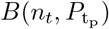 and 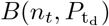, respectively. The number of migrating tumor cells is drawn from the respective binomial distribution 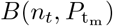 and then these cells are distributed around a neighborhood of 26 voxels (Moore neighborhood) according to a multinomial distribution [41]. For simplicity, it was assumed that all tumor cells belong to the same clonal population without including mutation events or phenotype changes.

For healthy cells, it was assumed that the levels of cell division and death remain balanced due to the ability of these cells to self-regulate. The biological process of migration is the only one that is affected by the evolution of tumor cells. Therefore, the number of migrating healthy cells is drawn from the binomial distribution 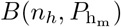 being displaced by the pressure exerted by the tumor cell colonization when the total number of cells in the voxel exceeds 45% of its maximum capacity. Figure 3 A shows a slice of an actual simulation, where the colors indicate voxel occupation. Note that each voxel can contain a different number of cells.

**Fig 3.**
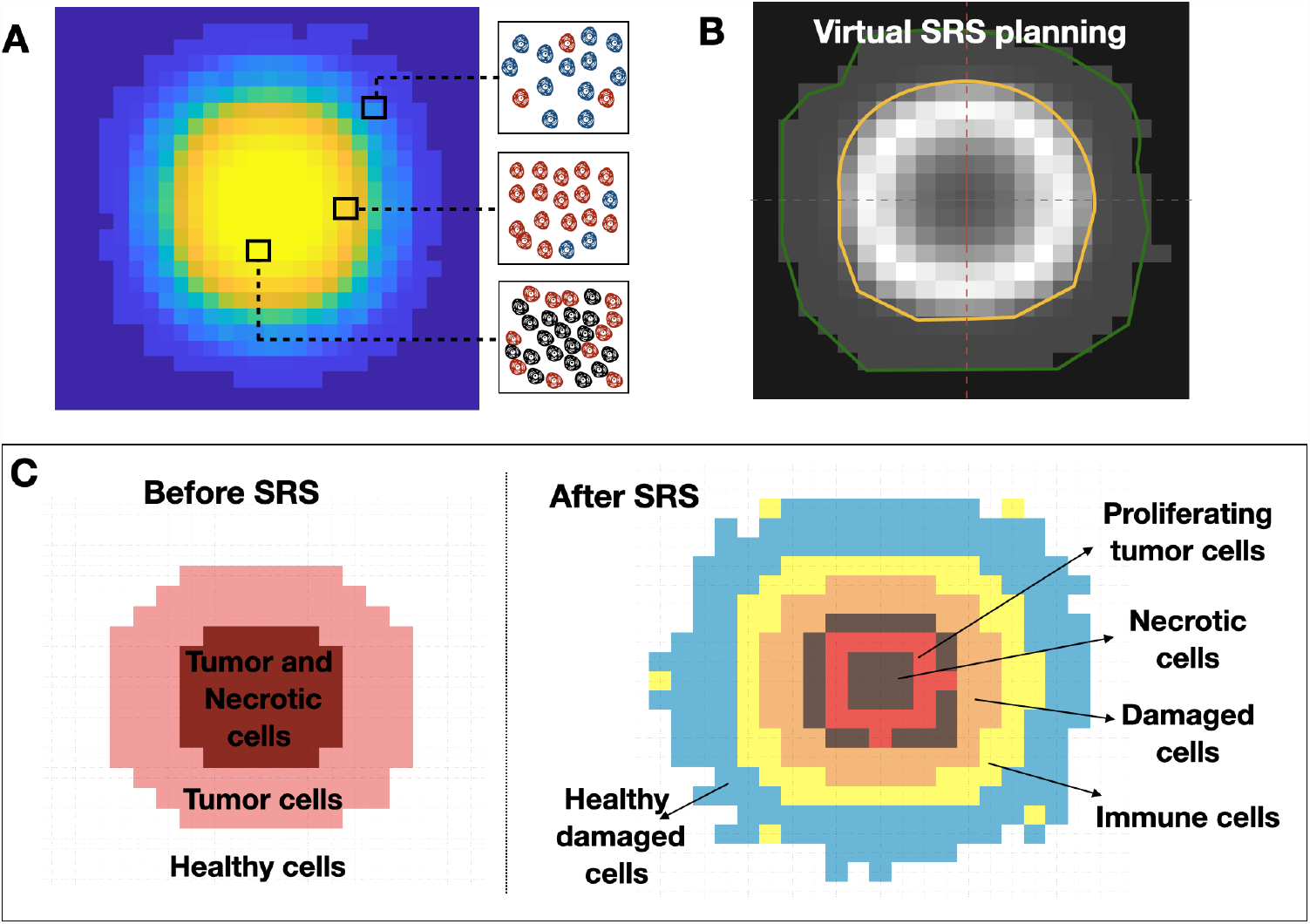
A Slice of a tumor simulation before SRS with DSBMS, mapping the distribution of cell density. A representation of the populations of healthy (blue), tumor (red) and necrotic (black) cells within a voxel according to their location is shown. **B**. Example of a single-shot treatment plan for a virtual simulation of SRS.The target is outlined in yellow, and it is the area most affected by SRS. The green line encloses another area affected with less intensity. **C**. Spatial distribution of cell populations just before and after SRS. Voxels may be occupied by more than one cell population but the dominant populations per voxel are shown.

To construct tumor growth rules in the DSBMS the probabilities of tumor cells reproduction, migration and death were first considered to be given by

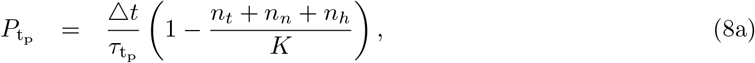

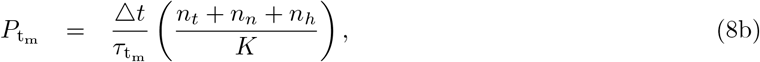

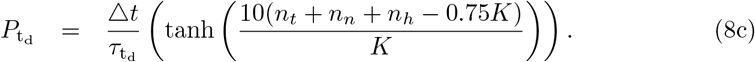

These probabilities were modulated by the relationship between the time step and the characteristic time of the process. The reproduction probability decreases with voxel occupation while the migration probability increases, simulating competition for space and resources. To mimic the effect of local vascular damage and hypoxia on tumor cell apoptosis signaling, it was activated once the voxel occupation reached a cell number greater than 75% of its carrying capacity. This was incorporated into the probability of death 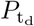 by the hyperbolic tangent term as a function of the voxel capacity.

As to motility, the probability of healty cells migration was given by

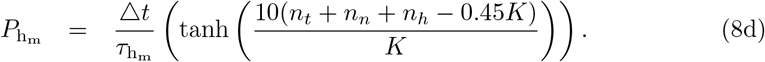

All probabilities were calculated at each time step and voxel, providing the number of cells undergoing mitosis, apoptosis and migration for every population. Parameter *τ* represents the characteristic times of each process. *K* is a carrying capacity.

#### Response to radiosurgery

A single dose of SRS was simulated in-silico. SRS leads to a fraction of tumor cells *S*_*f*_ surviving without damage and remaining viable, a fraction (1 − *S*_*f*_) receiving lethal damage. Of those, a fraction *ϵ* dies on a short time scale (i.e. days), and the remaining fraction 1 − *ϵ* moves into the compartment of damaged cells. Radiation therapy induces immune cells and lethal damage to a fraction *S*_*n*_ of healthy cells surrounding the tumor. Figure 3 C shows an example of the spatial distribution of cell population before and after SRS, as described above.

Let *n*_*d*_, *n*_*i*_ and *n*_*hd*_ be the total number of damaged tumor cells, activated immune cells and damaged healthy cells inside a given voxel, respectively. The numbers of damaged tumor and healthy cells that die by radiation are drawn from binomial distributions 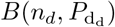 and 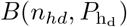, respectively. Then, the number of necrotic cells increases by adding these populations. Additionally, the immune system is activated and immune cells move to the irradiated region to remove necrotic cells. The number of necrotic cells eliminated by interaction with immune cells and the number of immune cells activated are drawn from binomial distributions 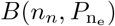 and 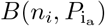, respectively. Furthermore, the activated immune cells are removed naturally and this process is simulated from the binomial distribution 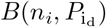. In an analogous way to tumor cells migration, the number of migrating immune cells is drawn from the binomial distribution 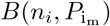 using the same algorithm.

Different probabilistic events where included to develop this part of the DSBMS. First, the probabilities of death of damaged cells are given by

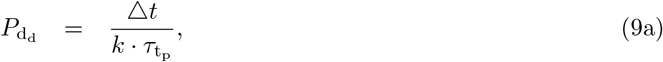

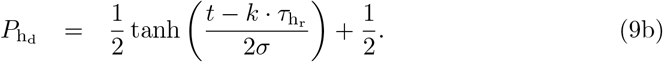

Damaged tumor cells die by mitotic catastrophe after *k* cycles of mitosis while trying to repair the damage caused [51]. The probability of this event (Eq. (9a)) is modulated by the relationship between the time step, the division time rate 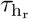 and the number *k* of the mitosis cycle. Similarly, damaged healthy cells die by mitotic catastrophe while trying to renew themselves. The time frame of this death event is longer since, as stated above, healthy cells have a low proliferation rate. For this reason, 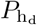 described by Eq. (9b) has a time dependence and a similar structure to the standard logistic function. This probability increases when damaged healthy cells are closer to their *k*-th division. Here *t* represents the time elapsed from radiosurgery to the evaluation step and 1/2*σ* is the compression parameter.

It was assumed that the probability of immune cells activation to be given by

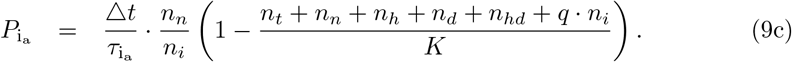

The immune system activation event takes into account the proportion of necrotic and immune cells within each voxel, and the characteristic activation time when there is at least one immune cell inside. 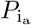 decreases with the voxel occupancy, simulating competition for space and resources. It was assumed that the size of immune cells is different from other cell populations, being *q* times the size of a healthy or tumor cell.

The probability of necrotic cells elimination is described in the DSBMS

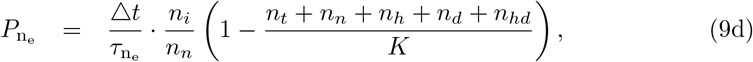

This probability was modulated by the ratio between the two populations, inversely to the activation process. In this case, 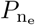 is also affected by the carrying capacity of the voxel.

Finally, the probabilities of immune cells death and migration were assume to have the form

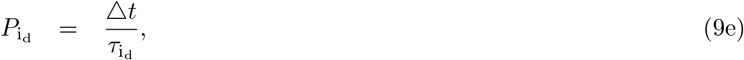

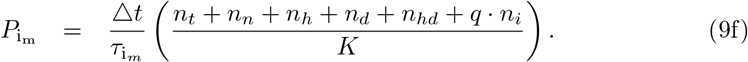

These probabilities were modulated by the relationship between the time step and the characteristic time of the process. Additionally, migration events were affected by voxel occupancy, making it harder to move in crowded contexts.

In all the previous formulations, the parameters *τ* included in each probabilistic event represent the characteristic times of each process (see Table 1).

**Table 1.** Summary of parameter values used for the stochastic model.

| Parameters | Meaning (average times per voxel) | Value (hours) |
| --- | --- | --- |
| $\tau_{t_p}$ | Tumor cells division time | 450-550 |
| $\tau_{t_d}$ | Tumor cells death time | 1000-1500 |
| $\tau_{t_m}$ | Tumor cells migration | 1000-2000 |
| $\tau_{h_r}$ | Healthy cells reproduction | 4680 |
| $\tau_{i_m}$ | Immune cells migration | 150-250 |
| $\tau_{i_d}$ | Immune cells death | 1440- 1560 |
| $\tau_{i_a}$ | Immune cells activation | 12-20 |
| $\tau_{n_e}$ | Necrotic cells elimination | 72- 96 |

#### Estimation of parameters for the DSBMS

We choose domain simulation sizes in line with those found in the clinical setting for metastatic lesions pre-SRS treatment, which are around 0.5 − 2 cm^3^ with maximum tumor sizes up to 10 cm^3^. Since high-resolution MRI voxel size is around 1 mm^3^, we chose that voxel size 1 mm^3^. Hence, space is discretized as a hexahedral mesh consisting of *L* × *L* × *L* spatial units (voxels), where *L* = 60. The time step was fixed to Δ*t* = 4 hours. From typical cell sizes [44], the 233 carrying capacity of a single voxel was estimated to be *K* = 2 × 10^5^ cells, assuming similar sizes for tumor and healthy cells (∼ 10-12 *μ*m). Basal rates of tumor cell division, death and migration were chosen using Bayesian criteria based on the doubling times estimations [45] and imaging data from real BMs in [26]. The value *k* = 2 was taken, i.e. damaged tumor cells dying by mitotic catastrophe after two cell cycles on average [51].

Parameters related to the immune system were obtained from data of the microglial cell population [46, 47], which constitutes 0.5%—16.6% of the total number of cells in the human brain and the most abundant type of immune cell in the brain. Then, the initial number of immune cells was set at 10% of the healthy cells surrounding the tumor [48]. Immune cells are activated if within voxels are cells damaged or necrotic by SRS. Furthermore, it is known that the immune cells increase in size by 50% (*q* = 3/2) after activation [49]. It was assumed that the mean lifetime of immune cells is around two months and the activation time is in the range of 12 to 20 hours [48].

All the proposed parameters are associated with cellular processes, which combined result in whole-tumor rates. Cellular traits were randomly sampled from the range of allowed basal rates for each simulation.

#### Virtual BMs simulations

To simulate tumor growth dynamics after SRS, a set of simulations of BMs was run, starting from 10^3^ tumor cells, allowing them to grow until they reached diagnostic volumes in the range 0.5 − 2 cm^3^. Radiosurgery events were simulated and post-treatment tumor evolution continued as previously described. Each simulation had a different set of basal rates, sampled randomly from the ranges specified in Table 1. This study focused its attention on cases with either late inflammation or progression, but always in cases with initial response. Thus, simulations displaying longitudinal volumetric growth in the first four months after SRS were rejected.

To simulate the effect of radiosurgery, the voxel survival fraction was defined

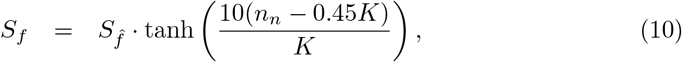

where 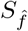 is the maximum survival fraction. The well-oxygenated cells are less resistant to radiation (more radio sensitive). Thus, cells that are farthest away and that do not get enough oxygen and nutrients to survive are those that are found in the voxels with the highest number of necrotic cells.

Two sets of simulations. were carried out. A first group of 200 BM simulations was performed under the condition of no damage to the healthy tissue surrounding the tumor (*S*_*n*_ = 1). This group would correspond to an idealized scenario, that is highly unlikely to happen in clinical practice. The second group of 200 BM simulations accounted for the damage induced by SRS to healthy tissue next to the lesion (0.1 ≤ *S*_*n*_ ≤ 0.7). The latter corresponded to the scenario that would be expected to happen in the clinical setting.

To calculate the lesion volume, the set of voxels reaching more than 45% of their carrying capacity was considered, looking at all cell types including necrotic cells. This set was denoted by *ν* and *N*_*ν*_ = |*ν*| was calculated as the number of elements in *ν*. Since the volume of a voxel was taken to be equal to 1 mm^3^, the tumor volume is equal to the number of occupied voxels *N*_*ν*_ in mm^3^.

#### Calculation of *β* exponent for virtual BMs

To study the dynamics of post-treatment volumetric growth for simulated tumors, the exponent *β* was calculated from the volumetric simulation data using Eq. (7).

To perform the calculation of *β* exponent for a virtual tumor, attention was paid to the dynamic behavior of tumors after the second follow-up (six months after radiosurgery). Taking into account that the average time between follow-ups in the clinic is three months, for each case, we calculated the following three volume measurements were calculated from six months post-SRS, that is, t = 6, 9 and 12 months. After that, the volumes were 282 fitted to the growth law and the value of the exponent *β* was calculated.

The three-time instances were taken within the following ranges: *t*_0_ ∈ (180, 180 + 15) days, *t*_1_ ∈ (*t*_0_ + 80, *t*_0_ + 100) days and *t*_2_ ∈ (*t*_1_ + 80, *t*_1_ + 100) days. Then, *β* was estimated for each simulation for the volume and combination of *t*_0_, *t*_1_ and *t*_2_. This procedure was repeated 20 times. Finally, we obtained the estimated 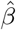 value was obtained for the corresponding simulated tumor as the median value of all the calculated values.

#### Computational implementation

The DSBMS was coded both in Matlab (R2020b, MathWorks, Inc., Natick, MA, USA) and Julia (version 1.5.3). The main workspace and simulation sections were coded in Julia, while data analysis and plotting were coded in Matlab. Simulations were performed on a 4-core 16 GB 2.7 GHz MacBook Pro. Computational cost per simulation was of the order of 1-3 minutes.

### Human data

The human data utilized in this study were sourced from the dataset provided in [40]. A dataset of BMs collected as part of a retrospective, multicenter, nonrandomized study, treated with stereotactic radiotherapy and followed up with magnetic resonance imaging (MRI). Cases with at least 3 consecutive imaging studies available, and classified as radiation necrosis, were used. The volumetric data used in this paper was calculated from volumetric contrast-enhanced T1-weighted MRI sequences, and the main interest was on total volume measurements.

## Results

### Compartmental mathematical model reproduces the accelerated growth dynamics of RN events

#### The compartmental model provides information on the evolution of cell populations and fits patient data

To simulate the RN event, we conducted simulations using the mathematical model described in Eqs. (1). This model allows us to track the evolution of each type of cell over time, providing insight into how the populations of different cell types change during the development of RN. Figure 4.a shows the results of one of those simulations. Initially, only tumor cells are present, and they grow exponentially. Meanwhile, necrotic cells steadily increase as damaged healthy cells die when they attempt to renew. Next, it was observed that the primary drivers of the fast-growing phenomenon associated with RN were the immune cell populations. These immune cells contributed the most to the total number of cells and played a critical role in the RN process.

**Fig 4.**
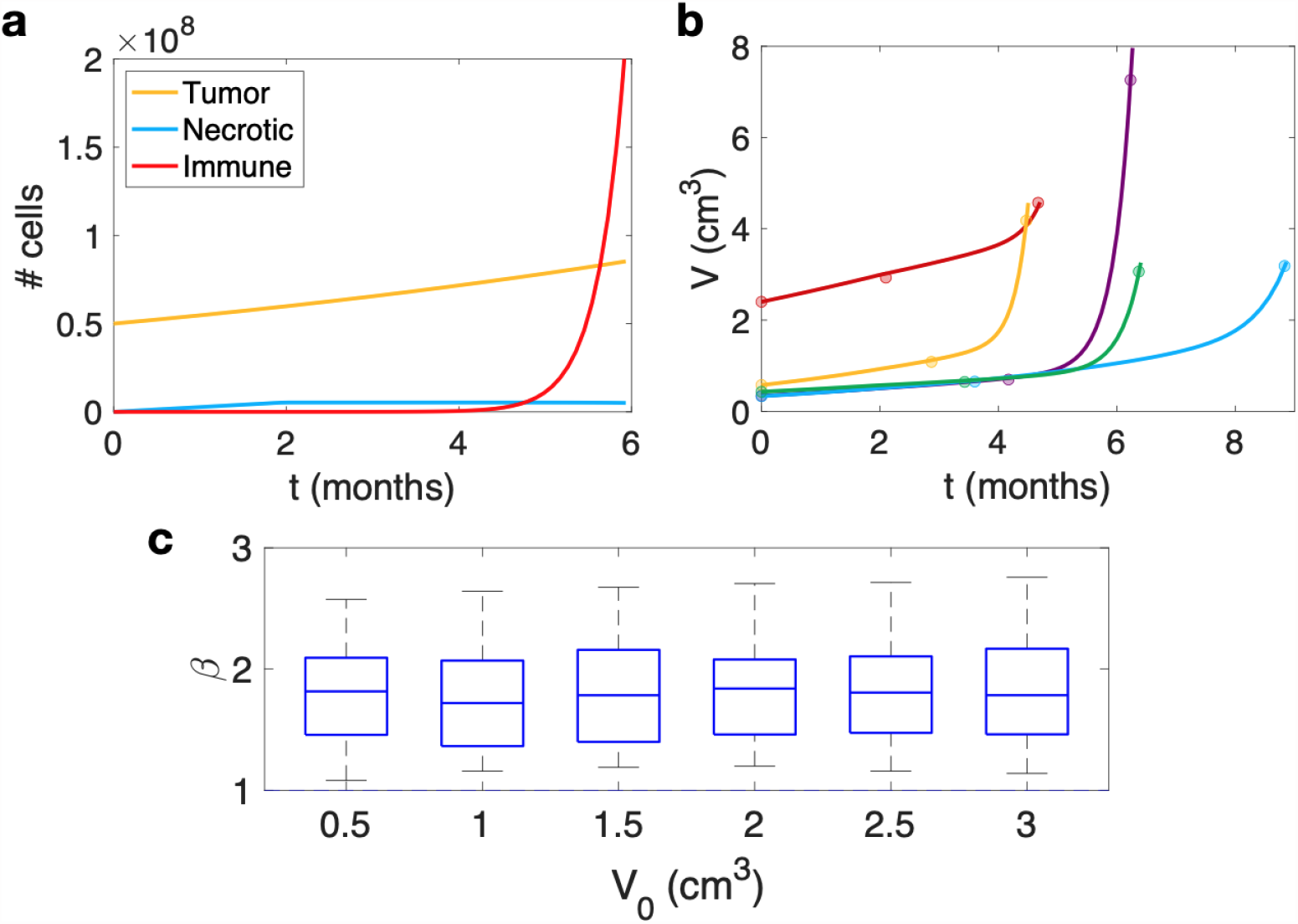
Results of simulations of the compartmental model (Eqs (1). **a**. Example of a simulation of model equations (1) showing the typical dynamics observed post-SRS in the context of RN events. The simulation illustrates the evolution of each cell type, with the initial number of tumor cells set to *N*_*T*_ = 5 · 10^7^, and parameter values *ρ* = 0.07 days^−1^, *λ*_*N*_ = 2.3 · 10^−11^ days^−1^, *γ* = 1.9 · 10^−7^ days^−1^ and *θ* = 0.17 days^−1^. **b**. Fittings using Eqs (1) (solid lines) for the longitudinal volumetric growth data (circles) for five patients diagnosed with RN. The parameters used for the fitting process are shown in Table 2. **c**. Distribution of growth exponents, *β* values obtained for simulations with varying initial volumes ranging from 0.5 to 3.0 cm^3^ by simulating 500 RN events per volume using a randomized approach where model parameters took values in the predefined range. Note that values of *β* between 1 and 2 were obtained systematically.

Fittings were also performed of the solutions of Eqs. (1) to actual data from patients diagnosed with RN. This led to finding individual parameter values providing excellent fits to the data. Figure 4.b shows five examples of these fitted lesions. (see Table 2 for the particular values for each fit).

**Table 2.** Model Eqs. (1) parameters best fitting the longitudinal dynamics of RN events. Volumetric data were taken from Ref. [40] and lesions 1 to 5 correspond to patients ID numbers: 30031, 40024, 40001, 40176, and 40042, respectively.

|  | Unit | #1 | #2 | #3 | #4 | #5 |
| --- | --- | --- | --- | --- | --- | --- |
| Initial volume, $V(0)$ | $\text{cm}^3$ | 2.4000 | 0.3400 | 0.4290 | 0.3312 | 0.5760 |
| Proliferative growth rate, $\rho$ | $\text{days}^{-1}$ | 0.0032 | 0.0057 | 0.0040 | 0.0063 | 0.0073 |
| Healthy cells dying, $k$ | $\text{days}^{-1}$ | $7.31 \cdot 10^5$ | $2.01 \cdot 10^5$ | $2.34 \cdot 10^5$ | $1.97 \cdot 10^5$ | $2.84 \cdot 10^5$ |
| Necrotic cells elimination rate, $\lambda_N$ | $\text{days}^{-1}$ | $2.14 \cdot 10^{-11}$ | $2.11 \cdot 10^{-11}$ | $2.27 \cdot 10^{-11}$ | $2.36 \cdot 10^{-11}$ | $1.85 \cdot 10^{-11}$ |
| Immune cells activation rate, $\gamma$ | $\text{days}^{-1}$ | $1.84 \cdot 10^{-7}$ | $1.99 \cdot 10^{-7}$ | $1.99 \cdot 10^{-7}$ | $2.06 \cdot 10^{-7}$ | $2.06 \cdot 10^{-7}$ |
| Immune growth rate, $\theta$ | $\text{days}^{-1}$ | 0.195 | 0.181 | 0.171 | 0.139 | 0.219 |
| Immune cells elimination rate, $\lambda_I$ | $\text{days}^{-1}$ | 0.07 | 0.07 | 0.07 | 0.07 | 0.07 |

#### Analysis of model behaviour reveals high growth exponents in terms of volume growth

To further explore the behavior of the mathematical model, we conducted simulations with different initial volumes ranging from 0.5 to 3.0 cm^3^. Specifically, we simulated 500 RN events per volume using a randomized approach where the model parameters took values within a given range. The exponents obtained in these simulations are shown in Figure 4.c. The findings revealed that the median *β* values obtained from the mathematical model are consistently larger than 1.5, regardless of the initial volumes or model parameters used in the simulations.

### Stochastic mesoscopic model reflects the observed dynamics in BM recurrences and RN events

The stochastic mesoscale brain metastasis growth simulator was set up using the biological rules described in ‘Methods’. A total of 400 simulations were performed of virtual BMs treated with SRS displaying growth after treatment and the exponent *β* was calculated. Figure 5 presents two simulation examples, capturing distinct scenarios of recurrence and radiation necrosis (RN) events, offering insights into their volumetric changes over time and the distribution of the diverse cell populations.

**Fig 5.**
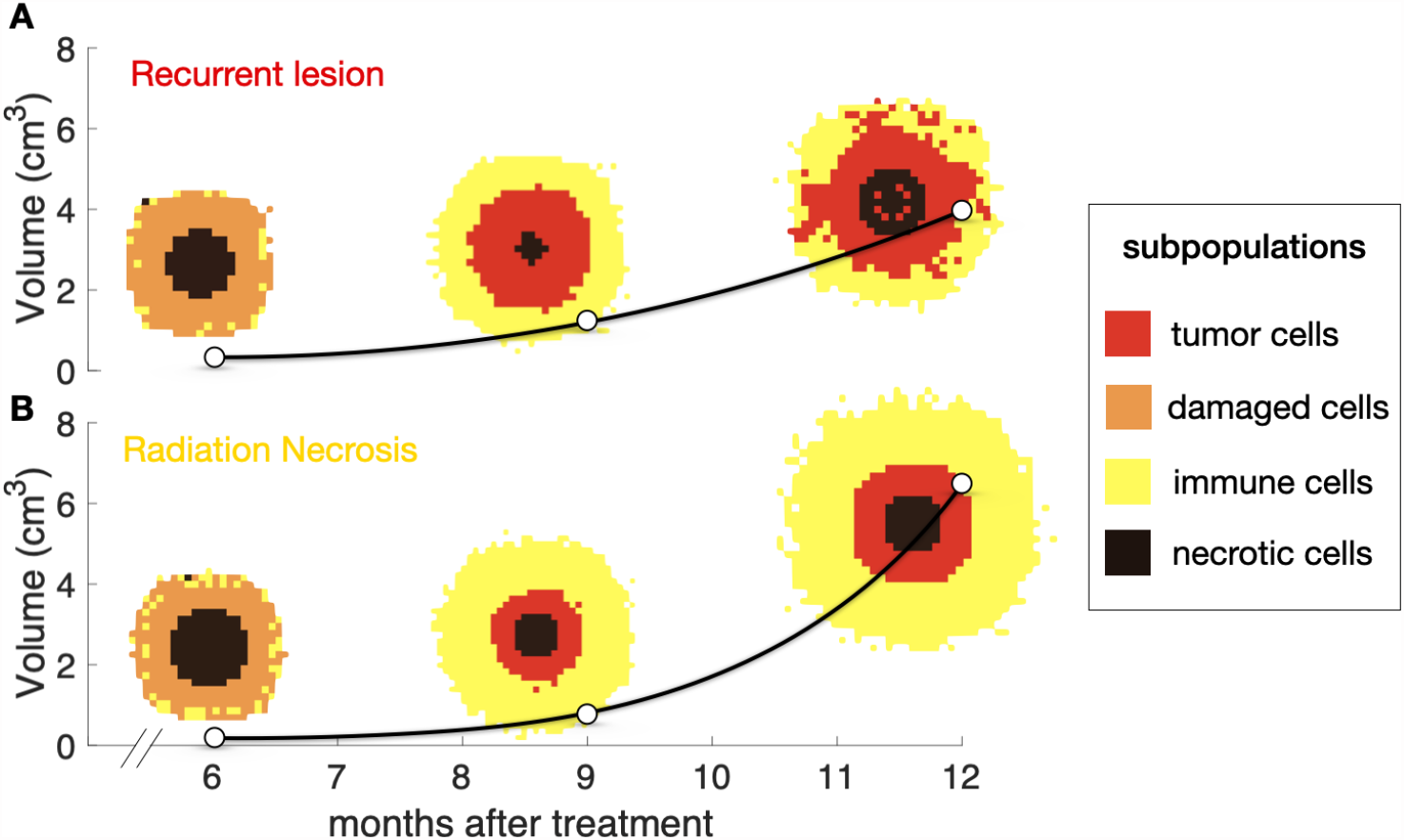
The dynamics of brain metastases (BMs) recurrences and radiation necrosis (RN) events are captured by the stochastic mesoscopic model. **a**. Recurrent BM. **b**. RN event. White points represent the total tumor volume at different time points, and black lines depict interpolations between the points (provided as visual guidance). Additionally, spatial distribution of the four cell populations illustrate the dynamic shifts in dominance among cell types for both scenarios. The dominant population per voxel is shown.

#### DSBMS correctly reproduces the volumetric dynamics of BM growth after SRS

The volumetric growth dynamics after therapy of the virtual BMs generated was 337 studied. Figure 6 shows three examples of these in silico simulations. The first column (A, C, E) shows the dynamics of the different cell populations present: proliferating tumor cells, damaged cells, necrotic cells, immune cells and total tumor cells for the three cases. The second column (B, D, F) shows the longitudinal volumetric dynamics of the simulation displayed in the first column. In each case, 20 *β* growth exponents were calculated as explained in ‘Methods’. Additionally, the median 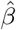 was obtained for each simulation. In two of the cases, sublinear growths 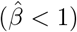 were obtained for recurrences. These simulations were generated with little or no damage to healthy tissue, i.e, *S*_*n*_ = 1 (Figure 6 (A,B)) and *S*_*n*_ = 0.7 (Figure 6 (E,F)). On the other hand, when there was substantial damage to healthy tissue *S*_*n*_ = 0.1, the volumetric evolution displayed a superlinear growth 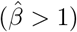, as shown in Figure 6 (C,D).

**Fig 6.**
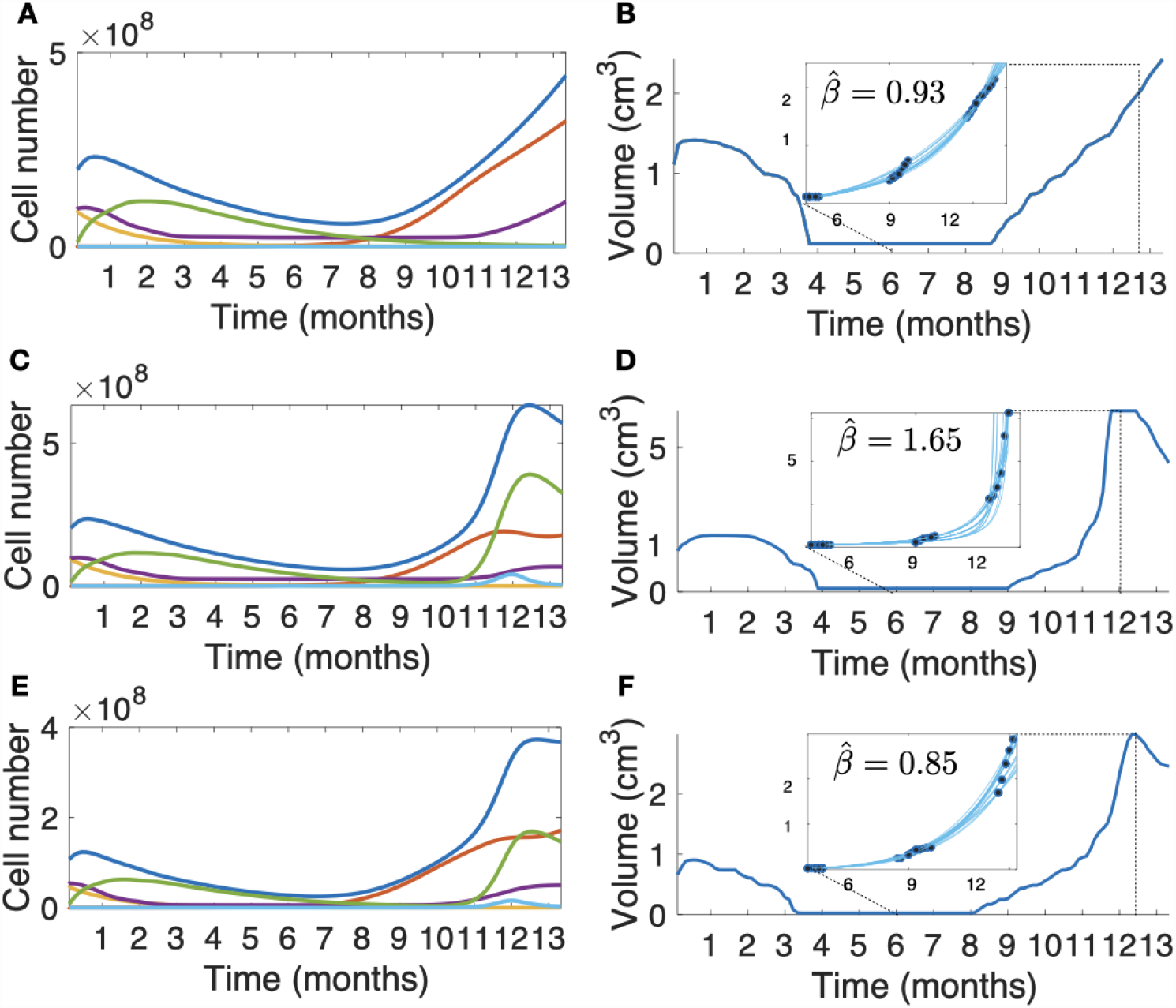
DSBMS simulations of longitudinal tumor growth dynamics after SRS. The first column (A,C,E) shows the dynamics of the total populations of proliferating cells (red lines), damaged cells (orange lines), necrotic cells (violet lines), immune cells (green lines) and total tumor cells (blue lines). The second column (B,D,F) shows the longitudinal tumor volumetric dynamics. Blue lines inside the zoom box represent different fits of *β* exponent solving Eq. (7) and the 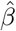 mean is shown. Subplots (**A-B**) correspond with a tumor simulation with no damage to healthy tissue 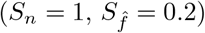, subplots (**E-F**) correspond with small damage to healthy tissue 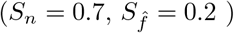 and subplots (**C-D**) correspond with high damage to healthy tissue 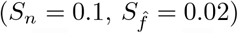. Basal rate parameters for simulations in this figure are 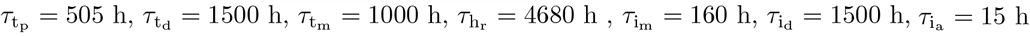 and 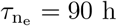.

#### Post-SRS BMs are characterized throught growth exponents

Thus, in order to characterize the volumetric growth post-SRS of the BMs using the scaling exponent, the values of *β* were calculated for the set of 400 virtual BMs. The results obtained for the first group (*S*_*n*_ = 1) are shown in Fig. 7 (A). The values of *β* were grouped according to the value of 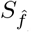 used in the simulation. Medians and quartiles of the box plots were mostly below 1, although for small values of 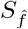, there were a set of outliers with high estimates of *β*. These exponents described the dynamics of relapsing lesions.

**Fig 7.**
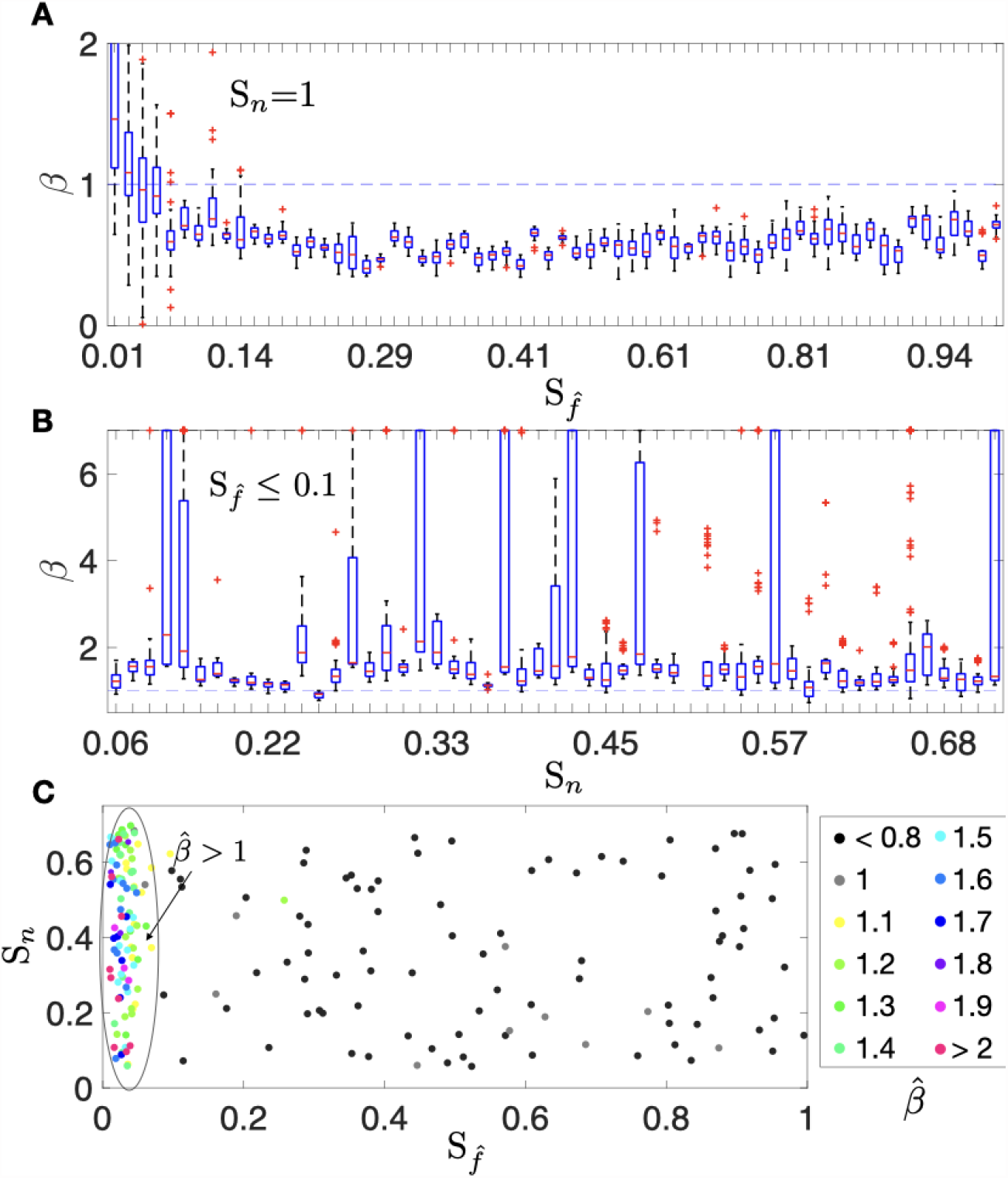
Comparison of box plots for the growth exponents *β* calculated for virtual BMs performed with the DSBMS. **A**. Growth exponents *β* values computed for the group of 200 simulations with *S*_*n*_ = 1 and 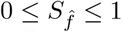. **B**. Growth exponents *β* values computed for the group of 200 simulations with 0.1≤ *S*_*n*_ ≤ 0.7 and 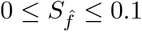. **C**. Scatter plot that shows the *β* median calculated for the virtual BMs which were simulated with different values of 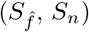. Basal rate parameters for the simulations are shown in Table 1.

For the second group, which included damage to healthy tissue, there were two behaviors observed in silico. Figure 7 (B) shows the scatter plot of the 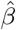 median calculated for the virtual BMs according to the different values of 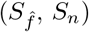 simulated. Values of 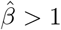 were obtained for cases where SRS eliminates most of the tumor cells (values of 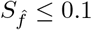). Here, the volume re-growth was due to the inflammatory component. Otherwise, for larger tumor remnants re-growth simulations (values of 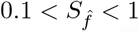) the calculated 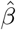 exponents were less than 1. This behavior corresponds to tumor relapse. In addition, the *β* values calculated for the cases simulated with 0.1 ≤ *S*_*n*_ ≤ 0.7 and 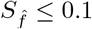 can be seen in Fig. 7 (C). Despite the observed variability, the *β* values obtained were typically greater than 1.

#### Growth exponents allow differentiation between radiation necrosis and recurrence

The computational results suggest that the *β* value could be used to distinguish the inflammatory response from tumor progression. The ANOVA test for the comparison with virtual BMs leads to significant differences between the inflammatory response group and relapses groups (p=1.85 × 10^−12^). Box plots for the different subgroups are shown in Fig. 8 (A). The area under the ROC curve (AUC) in Fig. 8 (B) illustrates the ability of the exponent 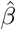 to discriminate between response groups. We obtained AUC=0.97 and the optimal threshold calculated to maximize the sensitivity and specificity values was *β*_*threshold*_ = 1.05. This means that inflammatory events show faster growth dynamics than relapses.

**Fig 8.**
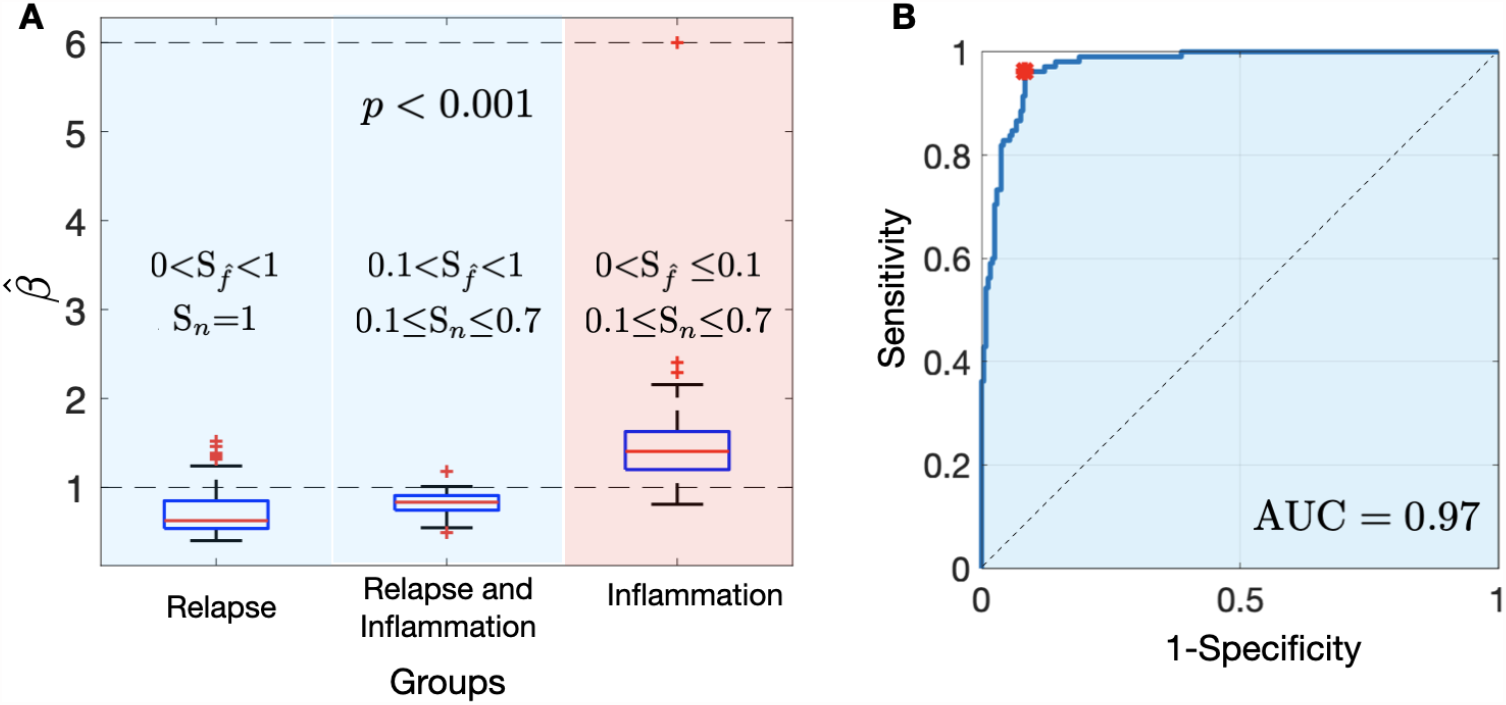
**A**. Box plots showing the comparison of the growth exponents 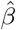 between the different simulated BMs: relapse group (R), whose response is characterized by tumor progression 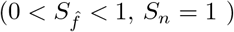, relapse and inflammation group (R & I), whose response is characterized by tumor progression and inflammation 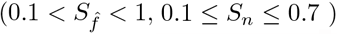 and inflammation group (I), whose response is characterized by inflammation 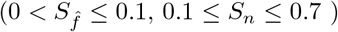. **B**. ROC curve for the discrimination between tumor progression (R and R&I groups) and inflammatory response (I group) according to the growth exponent 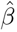.

## Discussion and Conclusion

Radiation necrosis is a common adverse effect associated with radiation treatment. To gain a deeper understanding of the growth dynamics of RN events, computational frameworks were developed, based on mathematical models of increasing complexity. These models provide mechanistic explanations for the observed growth dynamics of RN, and offer insight into the underlying mechanisms of this phenomenon.

Recent research has revealed that RNs tend to exhibit faster growth dynamics than recurrent BMs, which can help distinguish between the two entities. However, until now, no mechanistic explanation has been provided for these observations. The mathematical models developed in this work offer a promising way to fill this gap and provide a mechanistic understanding of the differences in growth dynamics between RN and BM.

Although RN is thought to be the result of synergy between vascular damage and immune activation, the precise mechanism underlying this phenomenon remains unclear [52, 53]. The first mathematical model presented in this study was a compartmental model. The hypothesis was that healthy tissue surrounding the lesion becomes damaged due to radiation, leading to necrosis and the activation of immune cells. This mechanism effectively reproduces the processes that take place during RN events. Additionally, the simulations consistently yield growth exponents larger than 1.5, regardless of the initial values.

However, there are limitations to this model. While it provides a basic understanding of the mechanisms involved in RN, it may be too simplistic for a more detailed understanding of the biological processes involved in the response to SRS. It is important to have a more comprehensive understanding of these processes, not only for RN but also for recurrent events. Differentiating RN from tumor progression is one of the main challenges in the management of BMs, since the treatment approach for each of these varies greatly, and accurate differentiation is essential for effective treatment [54]. An additional complication arises from the fact that RN is usually heterogeneous, where necrotic cells coexist with tumor cells [55]. This complicates the differentiation process and highlights the need for a more detailed understanding of the underlying biological processes.

To the authors’ knowledge, only two studies to date have implemented mathematical models to date to describe RN. The first study proposed a biomechanically coupled tumor growth model [31] that describes a logistic growth with a diffusive term in the presence of an external force. To estimate the model parameters, ten lesions were used, with five exhibiting tumor recurrence and five showing RN. The second study [32], based on the same model as the first, looked at 78 BMs. They assess the ability of the model to distinguish between tumor progression and RN, using not only a univariate analysis but also a radiomic analysis of 1080 radiomics features per patient, dividing the lesions into a training dataset comprising 66 BMs and a validation set of 12 lesions.

To account for the complexities of the system, a stochastic mesoscale simulator was also developed, providing a more detailed representation of the complex interactions occurring within BMs. In this simulator, BMs are described as a composition of different clonal populations seen at the voxel level, with three populations taken into account before treatment: healthy cells, tumor cells, and necrotic cells, and three more to account for the response after SRS: immune cells, damaged tumor cells, and damaged healthy cells.

Brain metastases growth was described on a mesoscopic scale in the simplest possible way, taking into account only one clonal population. Although it is known that these tumors present cellular heterogeneity in their composition, it is not fully characterized. For this reason, our results are not focused on describing the growth dynamics of BMs optimally. On the other hand, it has been known that tumor treatments induce a reduction of the clonal complexity at the point of maximal response due to the action of selective pressures of the drugs [50], which supports the simplification of a single clonal population to describe the response to SRS.

Continuous models can potentially neglect spatial correlations between the locations of individuals, especially when different species or subpopulations are taken into account. Therefore, the model developed here accounts for these relationships by combining discrete, spatial, and stochastic dynamics. This made it possible to analyze the dynamic behavior of the tumor in terms of volume through a more complete and precise approach. The model was used to simulate the different possible scenarios according to the damage caused to the healthy tissue surrounding the tumor, and the survival fraction of the tumor cells after therapy. Depending on whether the healthy tissue surrounding the lesion is damaged or not, this leads to different values of the growth exponents.

The discrete model explored the coexistence of necrotic cells with tumor cells, resulting in heterogeneous RNs. However, the interaction between the tumor and immune cell population was not taken into account, although it has been hypothesized that radiation induces damage signals within the tumor, making it for the immune system to detect [56]. There is only limited data available on the dynamics of immune response after radiation to a tumor lesion, but it could be interesting to include this in future work. Despite this, an increase in the necrotic cell population in the stochastic simulator indirectly affects the tumor cell population by saturating the voxels in question. In these cases, the probabilities of tumor cell proliferation and death are decreased and increased, respectively, and slower growth dynamics of this population are observed.

The stochastic simulator was tested to determine its effectiveness in distinguishing between recurrent lesions and RN events. The model classified both entities at 97% accuracy (AUC=0.97), which demonstrated its potential to differentiate between these two conditions. The threshold exponent *β* found, despite being larger than 1, was smaller than those computed by the continuous model and that observed in human beings. This can be explained by the inclusion of virtual simulations with small but not zero residual diseases (*S*_*f*_ *>* 0) after SRS in the inflammation group, which may affect overall volumetric growth dynamics. Another drawback is the low spatial resolution when cell densities decrease in response to therapy, which may affect the lesion volume measurement.

However, these results also suggest that the value of the exponent *β* could have a direct clinical application. The model supports that when three MRI studies are available satisfying the inclusion criteria, computing *β* for a particular patient could allow us to diagnose the growth as inflammation or tumor progression. Furthermore, this result is consistent with the study carried out in [34] on a group of real patients with and without radiation necrosis. In the present study, the area under the curve (AUC) associated with the receiver operating characteristic (ROC) curve was found to be 0.97, which is significantly higher than the AUC of 0.74 reported in the previously cited study that used actual patient data. One potential explanation for this difference could be the presence of confounding effects and a higher variability between patients in the actual patient data.

In conclusion, this paper has presented two models to explain the phenomenon of RN and to differentiate it from tumor progression. The first model, a simplistic continuous model, provides a plausible explanation for the high growth exponents observed in human beings during RN events. The second model, a stochastic mesoscale simulator, provided a more detailed representation of the complex interactions occurring within BMs, and explains the growth of BMs with varying degrees of damage. The simulator was tested to distinguish between recurrent lesions and RN events and proved to be a robust theoretical support to the use of the growth exponent *β* in differentiating between these two conditions. The development of accurate and detailed models could help improve the management of BMs and ultimately improve patient outcomes.

## Acknowledgments

This research has been supported by the Spanish Ministerio de Ciencia e Innovación (grants PID2019-110895RB-100 and PDC2022-133520-I00), Junta de Comunidades de Castilla-La Mancha (grant SBPLY/21/180501/000145) and BOT is supported by the Spanish Ministerio de Ciencia e Innovación (grant PRE2020-092178). The funders had no role in study design, data collection and analysis, decision to publish, or preparation of the manuscript.

## Notes

### Competing Interest Statement

The authors have declared no competing interest.

